# High-throughput phenotyping of single nucleotide variants by linking transcriptomes to genotypes in single cells

**DOI:** 10.1101/2023.05.22.541777

**Authors:** Sarah E. Cooper, Matthew A. Coelho, Magdalena E. Strauss, Aleksander M. Gontarczyk, Qianxin Wu, Mathew J. Garnett, John C. Marioni, Andrew R. Bassett

## Abstract

CRISPR screens with single-cell transcriptomic readouts are a valuable tool to understand the effect of genetic perturbations, but are currently limited because genotypes are inferred from the guide RNA identity. We have developed a technique that couples single-cell genotyping to transcriptomics of the same cells to enable screening for the effects of single nucleotide variants. Analysis of variants tiling across the *JAK1* gene demonstrates the importance of determining the precise genetic perturbation and classifies missense variants into three functional categories.

## Main text

Human genetics, population scale biobanks and cancer genome sequencing have identified thousands of genetic variants associated with disease^1,2^. However, the rate of discovery of such variants vastly exceeds our ability to understand and experimentally model their functional effects.

High-throughput CRISPR-mediated pooled screening for phenotype^3^ or coupled to single cell transcriptomics^4^ offers a powerful way to assess the effects of thousands of genetic perturbations. However, it is mainly limited to knockouts or manipulation of expression level using CRISPR interference or CRISPR activation since the guide RNA (gRNA) is used as a proxy of cell genotype and thus the efficiency of the perturbation must be very high. This makes it very challenging to screen for single nucleotide variants, since base editing, prime editing or homology-directed repair (HDR) efficiency is rarely high enough^5^, is highly variable between different genomic sites and cell types, and can lead to undesirable editing byproducts such as bystander mutations, insertions/deletions or heterozygous edits. Furthermore, even in those cases where base or prime editor screens have been successful^6,7^, we cannot distinguish cells containing a non-functional gRNA from those that have successfully introduced an edit without an effect, meaning that benign variants cannot be accurately classified.

To address these limitations, we developed a method, scSNPseq, that uses transcribed genetic barcodes to couple single-cell genotyping with transcriptomics to identify the exact genotype and transcriptome of each individual cell. This enables high-throughput pooled screening for SNVs with single-cell ‘omics’ readouts.

We used a previously described^8^ cytosine base editor screen in HT-29 cells with gRNAs tiling across the *JAK1* gene to establish our method. We have phenotypic data on the response of each variant to interferon gamma (IFN-γ) which triggers cell death and induction of PD-L1 and MHC-I expression, both of which are blocked by loss of JAK1 function^8^. Interrogated *JAK1* variants can inform the genetic basis of immunological disorders and mechanisms of cancer resistance to anti-tumour immunity.

Single-cell transcriptomics of base edited cells after IFN-γ treatment showed that cells fell into two broad clusters (Fig. 1a). To assign high-level identities to each cluster we assigned gRNAs to each cell (Supp Fig. 1a) and predicted the resulting edits (Supp Fig. 1d). We identified the two clusters as JAK1 loss of function (LoF) or not LoF by merging smaller clusters based on gene expression using the prevalence of cells with non-targeting gRNAs (NT-gRNA) in each cluster (Supp Fig. 1b, 1c). Stop codons and splice variants were predominantly contained in the LoF cluster, with WT, synonymous and intronic variants in the not LoF cluster (Fig. 1b, Supp Fig. 1e). This classification was confirmed by comparison with the results of previous screens for growth (proliferation score, Supp Fig. 1f) or induction of PD-L1 and MHC-I (FACS score) in the presence of IFN-γ (Supp Fig. 1g)^8^.

**Figure 1:**
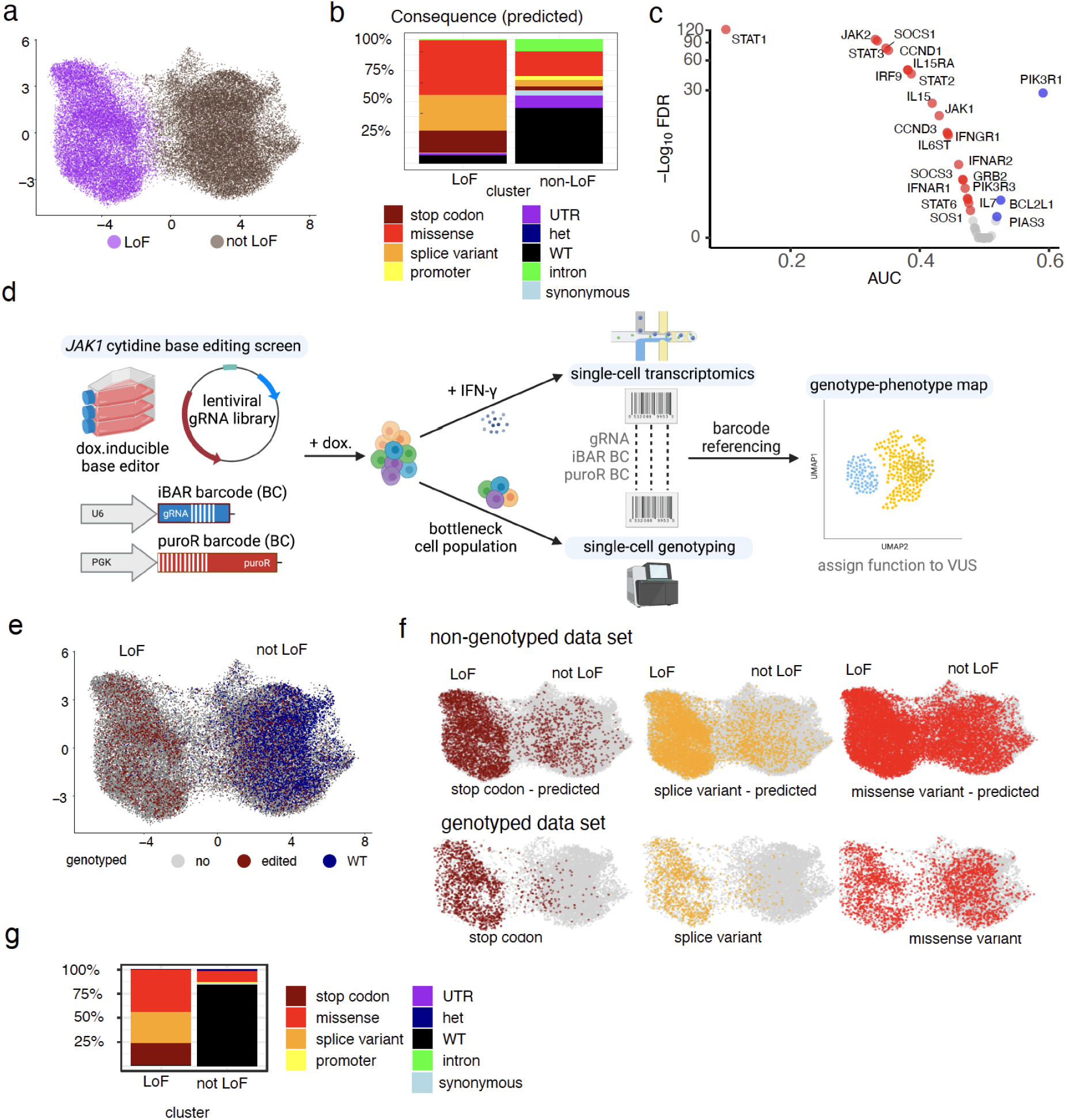
A single-cell base editor screen tiling across *JAK1* is improved by coupling genotype with transcriptome. a) UMAP of LoF and not LoF meta-clusters including all cells with a uniquely assigned gRNA. b) Distribution of consequences of the predicted mutations for each cluster. c) Differential gene expression analysis of JAK-STAT pathway genes between the LoF cluster and non-targeting gRNAs. AUC < 0.5 indicates downregulation (red, if significant), AUC> 0.5 upregulation (blue, if significant). d) Overview of high-throughput SNV phenotyping. Base editing of *JAK1* was achieved through the introduction of a barcoded gRNA library into doxycycline-inducible cytidine base editor expressing HT-29 cancer cells. After editing, cells were induced with IFN-γ before single-cell transcriptomics, or bottlenecked and processed for targeted single-cell DNA sequencing. Transciptomes and edited genotypes of single cells were linked through genetic barcodes to assign function to variants of unknown significance (VUS). e) UMAP combining the non-genotyped (grey) dataset with all genotyped cells with confidently called genotype (GT, 18,978 cells). Red and blue indicate edited and wild-type (WT) cells, respectively. f) UMAPs illustrating mutational consequences for the predicted genotypes compared to the actual ones. g) Percentages of cells showing the consequence of mutations from actual genotyping in LoF and not LoF clusters.

Analysis of differential gene expression between the two clusters showed a strong enrichment for components of the IFN-γ signalling pathway (Fig. 1c), including *JAK1* itself, *IFNGR1, JAK2, IRF9, STAT1, STAT2* and *STAT3* and downstream effectors such as *IL15, IL15R1, CCND1, CCND3*, and *SOCS3. STAT1* was one of the most downregulated transcripts in JAK1 LoF cells, suggesting a positive feedback loop may maintain *STAT1* mRNA expression in the presence of JAK1 signaling^9^. Also, the regulatory subunit of phosphoinositide-3-kinase (PIK3R1) was highly upregulated in the JAK1 LoF cells, consistent with extensive cross-talk between IFN-γ and PI3K signalling pathways^10^.

We next performed targeted single-cell genotyping to identify the precise mutations introduced in *JAK1* within each cell. To unambiguously couple the genotype to the transcriptome, the cells used for this screen had two different transcribed genetic barcodes introduced by lentivirus on the same vector as the gRNA library (Fig. 1d). One was in the 5’ untranslated region of the puromycin resistance gene (puroR BC), and a second within the first loop of the gRNA (iBAR BC). By using a targeted capture strategy, both barcodes can be read out in single-cell genotyping and transcriptomics assays (Methods). To accurately genotype the cells, we also bottlenecked the population to obtain multiple daughter cells from each edited cell, all of which are marked by the same barcode. We were able to confidently call the genotype of 233 barcodes with a minimum of 3 cells per barcode. These barcodes were represented by 18,978 cells in the transcriptomics analysis (average 81 cells/barcode) (Methods, Fig. 1e), which were used in subsequent analyses. For 25 gRNAs, we saw different barcodes for the same gRNA, resulting from multiple independent editing events (Supp Fig. 1h).

There was an improvement in classification of stop codon or splice variant mutations into the correct (LoF) cluster and WT cells into the not LoF cluster when considering actual genotypes (Fig. 1f, 1g), compared to using the gRNA as a proxy of genotype. Notably, missense mutations present in the not LoF cluster can be unambiguously defined as mutations that do not result in a loss of JAK1 function, rather than gRNAs that do not edit, and can therefore be used to assign these variants of unknown significance (VUS) as true benign mutations.

Transcriptomic changes resulting from the different mutations separated cells into two main clusters (Fig. 2a), those containing predominantly LoF mutations (stop codon, splice variant, some missense) or not LoF (WT, synonymous, some missense). We used diffusion maps^11^ to identify trajectories in the data (Fig. 2b), and the first diffusion component accurately reflected the trajectory between not LoF and LoF mutations (diffusion score, Methods). We confirmed this by comparison with JAK-STAT pathway activity (Supp Fig. 2a)^12^. The transcriptomic changes caused by the mutations split into two main clusters when ordered by diffusion score (Supp Fig. 2b) and correlated well with the differential expression of JAK-STAT pathway genes (Supp Fig. 2c). WT and synonymous variants had very low diffusion scores, stop codon or splice variants high diffusion scores and missense mutations were bimodally distributed between the two (Fig. 2c).

**Figure 2.**
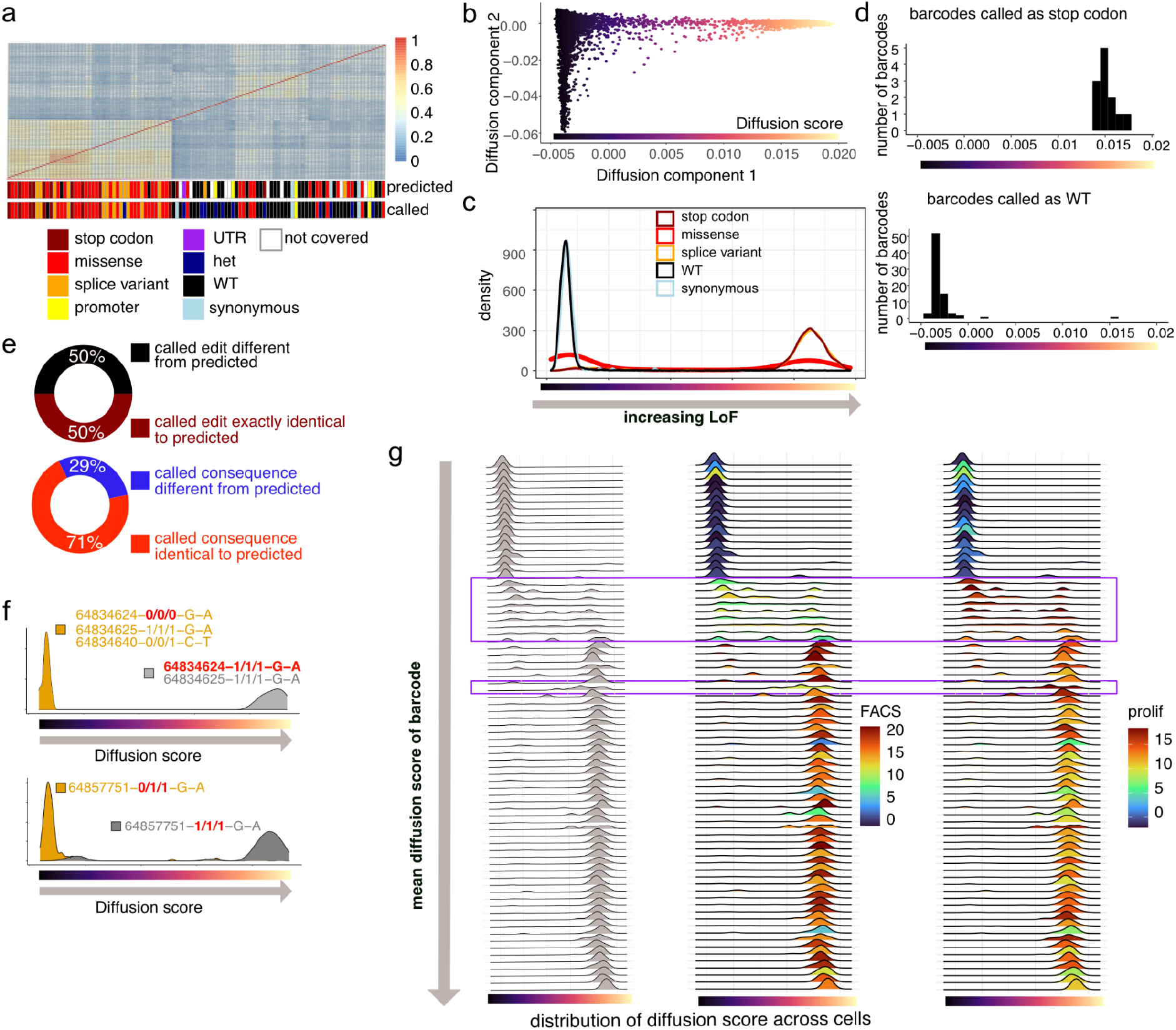
Transcriptomic changes of genotyped cells accurately classify missense mutations into three functional categories. **a)** Correlations of differential gene expression of each barcode to cells with WT-genotypes and non-targeting gRNAs. For each barcode, consequence and predicted consequence of homozygous mutations are shown. **b)** Diffusion map showing a low dimensional representation to identify the main directions of variation. **c)** The first diffusion component (diffusion score) acts as a measure of loss of function with high diffusion score for stop codons and low diffusion score for WT and synonymous mutations. **d)** Demonstration of low false negative and false positive rates for calling edits in barcode groups. Diffusion score for barcodes called as stop codons and WT. **e)** Percentage of barcodes with a predicted edit for which the called DNA editing is exactly the same as predicted based on complete editing in the window (maroon/black), and or for which the functional consequences of the edit are the same (red/blue). **f)** Possible phenotypic consequences of small differences in editing. Barcodes with the same gRNA, but different edits (heterozygous versus homozygous, one edit versus two consecutive edits). The position is the editing position on chr1. **g)** Transcriptomic heterogeneity of missense variants. Density plots for the diffusion scores of all barcodes with homozygous missense variants, including variants with low impact (low diffusion score indicating no LoF-benign), intermediate diffusion scores (indicating separation of function - SoF), and high impact (high-score missense) mutations. Boxed barcodes highlight variants with intermediate diffusion scores, characterised by lower FACS-scores and higher proliferation scores (SoF).

Cells with genotyped homozygous stop codon mutations were universally (100%, 12 out of 12) classified with high diffusion scores, and all 77 barcodes with WT genotypes except one (>98%) were classified with low diffusion scores (Fig. 2d). Out of the 15 barcodes called as homozygous splice variants, 93% (14) had high diffusion scores. Therefore, out of the 104 barcodes that were called with either a WT or a definite LoF genotype (stop/splice), 26 were true positives (definite LoF phenotype-high diffusion score), 1 was false positive (called as splice variant, but low diffusion score), one was false negative (precision: 96%, recall: 96%). This shows that our genotyping pipeline is highly effective and has a very low rate of incorrect genotype calls. This compares to 28 predicted stop/splice with 8 false positives, and 4 false negatives. (precision: 78%, recall: 88%) using the predicted genotypes.

When the actual genotypes were compared with those predicted from the gRNA sequence, only 50% of genotypes were exactly as predicted (Fig. 2e), although this was improved to 71% when analysed at the functional level due to degeneracy in codon usage (Fig. 2e). The incorrectly predicted cells contained a mixture of out-of-window editing, unedited cells, heterozygous edits, incomplete editing, and cells with different mutations on each allele (Supp Fig.2d, 2e, Fig. 2f). Looking at the position of the edits compared with the expected base editing window, 66% had an edit within the window, 25% contained no edit, and 9% contained an edit outside the window, most (8%) of which also had edits within the window (Supp Fig. 2f).

The benefit of genotyping is illustrated in two examples where we had the same gRNA associated with two different barcodes and where the genotype of these barcodes was different (Fig. 2f). In the first, both barcodes had a homozygous edit at chromosome 1 position 64834625, but only the clone that was additionally edited at position 64834624 showed a LoF phenotype, indicating that this mutation or the combination of the two together was causing the loss of JAK1 function. In the second example, only the homozygous edit at position 64857751 showed a LoF phenotype, whereas the heterozygous edit did not. Taken together, these observations demonstrate the utility of genotyping editing events to unambiguously interpret variant function, even in a screen optimised for very high base editing activity.

Some of the missense mutations had a diffusion score between the WT and LoF values, suggesting an intermediate phenotype (Fig. 2c, Fig. 2g). In our previous screen, these gRNAs had strong effects in the proliferation assay (prolif.), but weaker effects on PD-L1 and MHC-I protein expression (FACS, Fig. 2g), suggesting they could be separation of function (SoF) variants^8^. Closer analysis revealed that cells with these cell barcodes (and thus deriving from the same cell clone) were distributed across the diffusion score range. This shows that for these variants there is a stochastic response to IFN-γ, with some cells responding as normal, others not at all, and some with an intermediate effect. This may help to explain the difference between their long term effects on cell growth (prolif, Fig 2g) and their immediate effects on protein expression (FACS, Fig 2g), since growth integrates across time, whereas protein expression is a snapshot of their immediate response. SoF variants showed differential expression of IRF9, a key regulator of IFN-γ signalling, that may control the threshold of transcriptional response between WT, SoF and LoF (Supp Fig. 2g). These observations would not be possible without genotyping and single cell analysis.

In summary, we present scSNPseq, a technique that allows high-throughput pooled screening of single nucleotide or other defined variants with a single-cell transcriptomics readout through coupling of accurate single-cell genotyping and transcriptomics. We demonstrate its effectiveness in a base editor mutagenesis screen across *JAK1* to classify LoF missense variants, but importantly to identify benign variants, or variants with an intermediate phenotype (Supplementary Table 1). The methodology is applicable to any other methods for introducing variation such as HDR, prime editing^13^ or saturation genome editing^14^ since it does not rely on gRNA identity to call genotype. Due to the single-cell readout, it can be applied in a cell-type and state-specific manner, and to primary cells where the inability to clone cells normally prevents analysis of SNVs. It also provides a rich phenotypic readout of the whole transcriptome for each perturbation allowing classification of variants based on transcriptional signatures, and comparison to perturbations in disease. We believe scSNPseq will be invaluable for screening the functional significance and downstream effects of the growing list of coding and non-coding variants identified from human genetics analyses such as GWAS and cancer genome sequencing.

## Methods

### gRNA library cloning to include PuroR barcode and iBAR barcode libraries

To introduce the PuroR barcode (in the 5’ UTR of the puromycin resistance gene) a single stranded ultramer containing NeoUTR3^15^ was amplified using KAPA to add Gibson arms and a 12N barcode in the reverse primer. After SPRI purification, the product was cloned using Gibson assembly into lentivector (Addgene #67974) cut with XbaI and XhoI. After ethanol precipitation, 5 Gibson reactions were electroporated into supercompetent cells (Endura, Lucigen) and grown in liquid culture to give a coverage of around 100 million barcodes. gRNA with iBAR barcodes were introduced into the PuroR library by amplifying the gRNA library tiling *JAK1*^*8*^ (Twist, 2,000 guides) to include a 6N iBAR barcode in the primer. After a nested PCR, the gRNA iBAR library was cloned by Gibson into the PuroR library cut with BbsI and BamHI. After ethanol precipitation, 2 Gibson reactions were transformed into supercompetent cells and grown to give a coverage of around 40 million events. All primers are detailed in Supplementary Table 2.

### Base editing screens

For base editing experiments, we derived a clonal line of HT-29 cells expressing a base editor (cytidine BE3-NGG) under a doxycycline-inducible promoter^8^ and introduced the lentiviral gRNA library tiling *JAK1* with PuroR and iBAR barcodes as described above. We used an infection rate of ∼30% to minimise the introduction of multiple gRNAs in one cell, and selected infected cells with 2 ug/ml puromycin (Thermo Fisher Scientific). Cells were maintained in 0.5 ug/ml puromycin for the duration of the experiment to maintain gRNA expression. Base editing was induced by addition of doxycycline (1 μg/ml; Sigma Aldrich) for 72 h. After editing, we bottlenecked a subset of these edited cells (15,000 cells) and also used FACS^8^ to select LoF (50,000 cells) to ensure we captured representative phenotypes in our bottlenecked populations. After expansion, these cells were both loaded onto the Chromium X (4 lanes, aiming to recover 60,000 cells per lane) for transcriptomic experiments (see below for further details) and were also further bottlenecked (8,000 cells) for the genotyping plus transcriptomic experiments. After further expansion, these cells were single-cell genotyped with the Tapestri machine (Mission Bio, according to manufacturer’s instructions), using 4 reactions, up to 10,000 cells per reaction and using a custom panel of amplicon sequences (Supplementary Table 2) spanning *JAK1* exons and promoter region, as well as the gRNA plus iBAR barcodes and PuroR barcodes. The same population of cells were also loaded onto the Chromium X (2 lanes, aiming to recover 60,000 cells per lane). For all transcriptomics experiments, the base editor was induced again for 24 h as we have found it necessary to have expression of Cas9 in order to stabilise the gRNA transcripts and improve gRNA detection in single cells. We stimulated cells with IFN-γ (400 U/ml; Thermo Fisher Scientific) for 16 h before processing cells. We used the 5’HT kit (10X Genomics) and cDNA libraries were prepared according to manufacturer’s instructions. We performed direct gRNA capture by spiking in a scaffold specific RT primer before loading, and after the cDNA amplification we performed a nested PCR from the small SPRI fraction to produce a library for sequencing both the gRNA and the iBAR barcode. We also spiked in a puromycin resistance gene specific RT primer, and carried out an analogous nested PCR in order to produce a PuroR barcode library (primer sequences in Supplementary Table 2). Sequencing was performed on the NovaSeq 6000 (Illumina).

### Data analysis of single-cell base editor screen without genotyping

#### Processing and quality control

We used Cell Ranger 7.0.1 to obtain UMI counts for gRNA and mRNA and for cell-calling. For quality control, we removed low outliers for the total count, low outliers for the number of detected features and high outliers for the percentage of counts from mitochondrial genes using the scater^16^ Bioconductor package, obtaining 155,429 cells.

#### gRNA calling

We developed a robust method to call gRNAs and other barcodes in cells from (UMI) counts using a probabilistic model of mixtures of skewed normal distributions with three components. We considered all UMI counts above a minimum threshold of 2 in all cells. Then we used the mixture model to group them into three clusters, one cluster for ambient background noise, and two clusters for signal counts, to allow for a bimodal distribution of signal counts. For robust gRNA assignment and to exclude undetected multiple gRNA assignments in a cell, we defined two thresholds for UMI counts; a lower threshold – UMI counts below this threshold mean a 90% probability of being in the ambient cluster; and an upper threshold-UMI counts below this threshold correspond to a 10% probability of being in the ambient cluster. A gRNA was then called in a cell if UMI counts for 1 gRNA are above the upper threshold and no other gRNA has UMI counts above the lower threshold. We obtained 43,639 cells from this robust assignment of one gRNA per cell, which we used for downstream analysis. Using only cell barcodes with one gRNA assigned to them also removed most doublets, as these would have two gRNAs.

#### Dimensionality reduction and clustering

First, genes that are differentially expressed (DE) for at least one gRNA (with at least 10 cells assigned to it) compared to cells with non-targeting gRNAs are identified using the Wilcoxon rank-sum test^2^. Then we performed principal component analysis (PCA) on the data, subset to the DE genes and the genes in the JAK-STAT pathway. Louvain clustering^3^ was performed on a neighbourhood graph using 10 nearest neighbours for each cell, based on the low dimensional representation obtained by the PCA (Supp Fig. 1a). Two larger meta-clusters (referred to as WT (wild-type) and LoF (loss-of-function) are formed by grouping clusters by the percentage of cells with non-targeting gRNAs in the cluster (Supp Fig. 1b, Fig1a).

#### Differential expression analysis for LoF gRNAs

gRNAs for which at least 70% and at least 3 cells are in the LoF cluster were assigned to the LoF group. Differential analysis was performed between all cells of the LoF group and all cells with non-targeting gRNAs using the Wilcoxon rank-sum test (Fig. 1c). The Wilcoxon rank-sum test is a standard non-parametric test that compares for each gene how often its expression is higher for the LoF group compared to the cells with non-targeting gRNAs. Genes more highly or lowly expressed significantly often at FDR level of 0.1 are highlighted in Fig. 1c. The AUC (area under the curve) is the proportion of times that the expression of a gene is higher for the LoF group than in a corresponding cell of the non-targeting group, where corresponding refers to being the same quantile within the respective group. Therefore, AUC < 0.5 means downregulation in the LoF group, and AUC>0.5 upregulation. Using a non-parametric approach like AUC is more appropriate and robust for cases where a set of cells cannot be assumed to follow a parametric distribution like a Gaussian or a negative Binomial distribution. Here, we cannot assume cells of the same barcode that have been perturbed to follow the parametric distribution, as the cells may have been impacted to different degrees. An extreme example of this are the SoF mutants (Fig. 2g).

### Experiment with genotyping: analysis of scDNA-seq modality

The Tapestri DNA Pipeline On-prem was used for QC, cell barcode correction, alignment and cell calling, using as the reference the hg38 genome with pKLV2 added (Supplementary Table 2). For each cell, variant calling was performed using GATK HaplotypeCaller^17^. gRNA, iBAR, and puroR counts were computed for each cell barcode, using the reads for pKLV2 from the aligned bam files. Then gRNAs, iBARs and puroRs were assigned to cells using the same gRNA calling method as described above for the scRNA-seq modality. Groups of cells from the same parent cell (barcode groups) were identified as groups that either share the same gRNA-iBAR combination and the same puroR. For cases where either of the barcodes could not be called in a cell (because of noise), the assignment to groups was performed on the basis of the barcode called (iBAR or puroR). We obtained 490 unique barcodes with at least 3 cells.

Genotypes were then called on a per barcode-group basis, to allow robust genotyping for single-cell data, which have higher noise levels than pooled data and may be affected by allele dropout as well as distortion of genotype calling because of ambient counts. First, we subsetted cell genotypes to C->T and A->G mutations (for gRNAs on the reverse strand), and removed frequent mutations occurring in more than 10% of the barcodes, as we assumed that they were not caused by the gRNAs. We called genotypes for barcode-groups with at least 3 cells. For each position in the genome a variant was called if it was present on at least one allele in at least 2 cells from the group comprising at least 50% of the cells, and if a majority of cells with the variant have this variant on the same number of alleles. This relatively low threshold of 50% reflects the fact that it is unlikely that more than 2 cells and more than 50% of the cells of a barcode-group have a miscalled mutation by chance, and limits the impact of dropout and missed mutations on genotype calling at the level of barcode-groups. A barcode-group was called as WT, if for each position no more than 1 cell (or 0 cells if <10 cells per barcode-group) have a mutation on any number of alleles. The accuracy of this approach of genotype calling at the barcode-group level is shown in Fig. 2d. At this level of robustness and accuracy, we were able to call genotypes for 233 barcodes (Fig 1e, Supplementary Table 1), out of 490 barcodes with at least 3 cells identified overall (48%).

Consequences were assigned to edits on the barcode-group level using VEP^18^, restricting to MANE select proteins. Edits in the *JAK1* promoter region (chr1:64964978-64967543)^8^ were labelled as promoter. For several edits for a genotype we call the most severe consequence, where stop codon/start lost > splice variant > missense variant > promoter/intron > synonymous. Detailed genotype calls per barcode with consequence and additional analysis results can be found in Supplementary Table 1.

### Experiment with genotyping: analysis of scRNA-seq modality

#### Basic processing and gRNA calling

Basic processing and gRNA calling was performed in the same way as for the non-genotyped data. iBAR and puroR calling was performed as follows: first, a list of all possible iBARs was created and a list of puroRs was obtained from the puroR calling at the scDNA level. These lists were used as input in the cellranger pipeline, to obtain UMI counts for iBARs and puroRs in the same way as for gRNAs. Finally, iBARs and puroRs were called in cells using the same method as for gRNAs. Dimensionality reduction was also performed in the same way as for the non-genotyped data set.

#### Integration with non-genotyped dataset

To compare the genotyped to the larger non-genotyped dataset at the level of UMAPs and clusters, we used mutual nearest neighbours^19^ for data integration, and, based on the integrated PCA representation, assigned to each cell in the genotyped data set the UMAP coordinates of its nearest neighbour in the non-genotyped data set (Fig. 1e), and the most frequent cluster among its 10 nearest neighbours in the non-genotyped data set (Fig. 1g). For the clusters in Fig. 1g, a cell was filtered out if it was the only cell with a specific barcode within a cluster, to denoise possible errors in barcode assignment for the scRNA-seq data.

#### Correlation of differential expression across barcodes

Fig. 2a shows correlations of differential gene expression of each barcode to cells with both WT-genotypes and non-targeting gRNAs. The differential expression compared to the non-targeting cells with WT-genotypes was computed for each gene and each barcode with at least 10 cells. Then we computed the correlation across the AUCs obtained by this differential expression analysis, including for the computation of the correlation genes significantly differentially expressed for at least one barcode.

#### Diffusion and pathway scores

Diffusion maps^11^ were used to identify trajectories in the data. The first diffusion component, which we identified as the trajectory towards full LoF of JAK1, was named diffusion score. The pathway score for the JAK-STAT pathway (Supp. Fig. 3a) was computed using the PROGENy tool^12^.

#### Estimation of false negative and false positive genotype calls

We estimated the accuracy of our computational approach to genotyping at the barcode level using stop codons (which we can assume to lead to LoF) and WT (which cannot be LoF). We estimated the number of false positive genotype calls by examining the number of barcodes called as stop codons or splice variants, but with a diffusion score indicative of not LoF. Similarly, false negatives were estimated by considering the number of barcodes called as WT, but with a LoF phenotype (Fig. 2d). False positives and negatives for predicted rather than actually called phenotypes were estimated using predicted genotypes, excluding those gRNAs targeting the JAK1 promoter or UTR region and not covered by an amplicon.

#### Characterisation of SoF variants

We explored heterogeneity of LoF level of homozygous missense variants by means of density plots for the diffusion scores of all barcodes with missense variants, including variants with low impact (low diffusion score indicating no LoF-benign), intermediate diffusion scores (indicating SoF), and high impact (high-score missense) mutations (Fig. 2g). The plots (one density plot for each barcode) are ordered vertically by the mean diffusion score across the cells with the barcode. Barcodes with intermediate diffusion scores are highlighted by a purple box. A second, smaller, purple box highlights one additional barcode, to illustrate that this barcode has the same genotype as one of the barcodes in the first box. The variants highlighted by the boxes are characterised by lower FACS-scores and higher proliferation scores (SoF).

Specific gene regulation differences between SoF and full-impact missense mutations were identified as those either upregulated significantly for SoF compared to full-impact and not downregulated for SoF compared to benign missense variants (AUC>0.45), or downregulated significantly for SoF compared to full-impact and not upregulated for SoF compared to benign missense variants (AUC<0.55, Supp Fig. 2g). These cutoffs distinguish these genes from those that are upregulated compared to high-score missense mutations and downregulated compared to benign missense mutations, i.e. their gene expression is on a progressive trajectory between benign and full LoF (area highlighted in yellow in Supp Fig. 2g).

#### Data Availability

Sequencing data are deposited in the European Nucleotide Archive (ENA) with the accession ERP133355.

Code is available on GitHub (https://github.com/MarioniLab/scSNPseq) and processed data files on Zenodo (10.5281/zenodo.7941286).

## Acknowledgments

This research was funded by the Wellcome Trust Grant 206194 and Open Targets (OTAR2061). M.S. is supported by the Wellcome Trust (220442/Z/20/Z). Schematics were created with BioRender.com. For the purpose of Open Access, the author has applied a CC BY public copyright licence to any Author Accepted Manuscript version arising from this submission.

## Author Contributions

S.E.C., A.R.B and M.A.C. conceived the project. M.E.S. and Q.W. performed computational analysis. S.E.C., M.A.C. and A.M.G. performed wet-lab experiments. A.R.B., M.J.G. and J.C.M. supervised the project. S.E.C., M.A.C., M.E.S., A.R.B. drafted the manuscript with contributions from other authors.

## Supplementary information

**Supplementary Figure 1.**
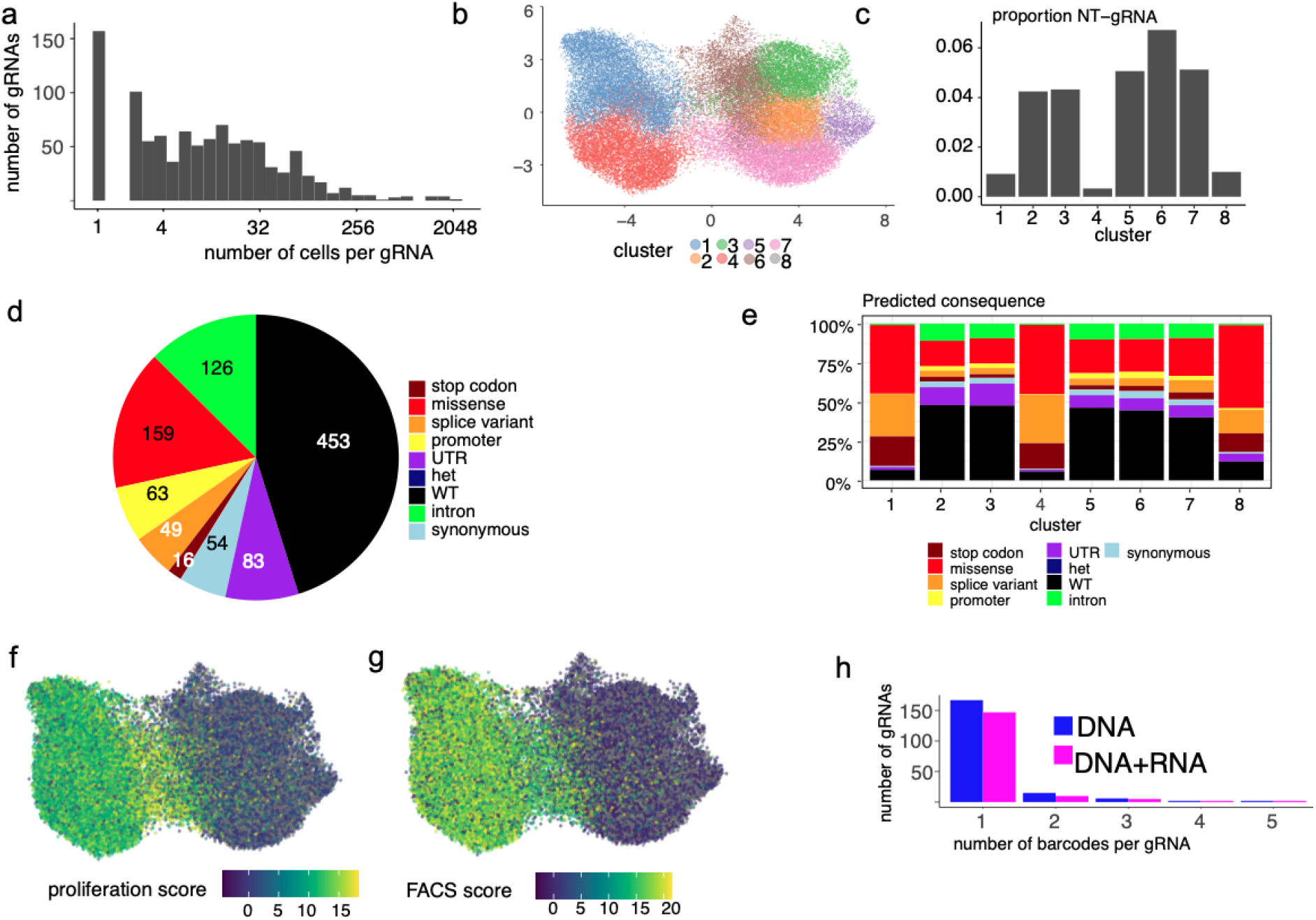
A single-cell base editor screen tiling across *JAK1* is improved by coupling genotype with transcriptome. a) Distribution of numbers of cells per gRNA for the experiment without genotyping. Overall, there are 1,003 gRNAs represented with at least one cell, and the mean number of cells per gRNA is 44. b) Clusters found by clustering the 43,639 cells with a unique gRNA assigned confidently using Louvain clustering on a neighbourhood graph using 10 nearest neighbours for each cell. c) Proportion of cells with non-targeting gRNAs groups clusters into two meta-clusters (1-4-8; 2-3-5-6-7). d) Number of gRNA for each predicted consequence for the non-genotyped experiment. e) Distribution of predicted consequence across clusters. A cell is assigned the predicted consequence based on complete editing within the editing window of the gRNA. f) UMAP highlighting proliferation score^8^. g) UMAP highlighting FACS-score^8^. h) Number of genotyped puroR/iBAR barcodes per gRNA for the genotyped experiment. Barcodes also present in the RNA modality are shown in magenta, those only present for the DNA in blue.

**Supplementary Figure 2.**
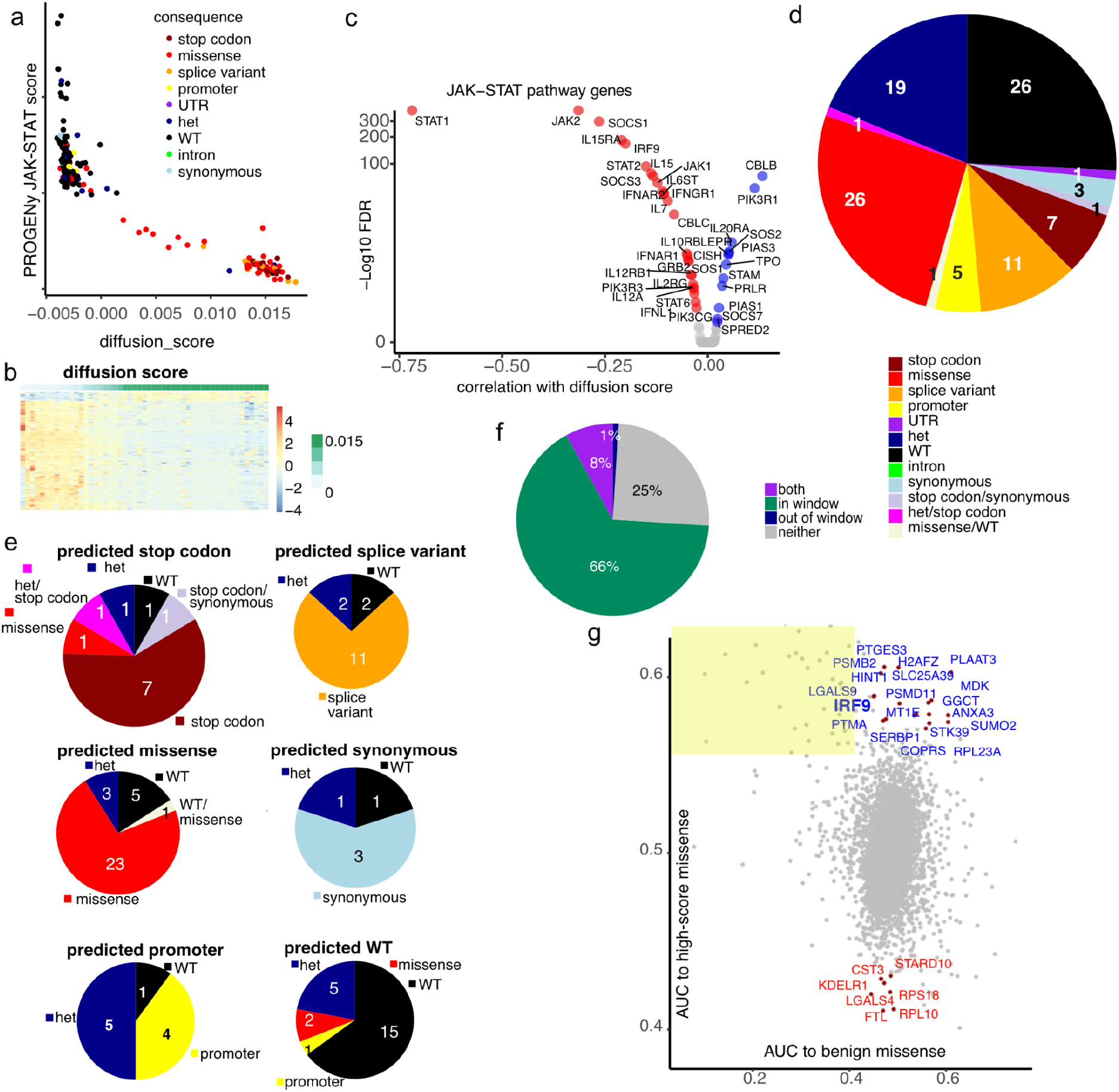
Transcriptomic changes of genotyped cells accurately classify three categories of missense mutations into three functional categories. a) JAK-STAT pathway activity, measured by the PROGENy score, is negatively correlated with the diffusion score. Each dot is a barcode, coloured by consequence. PROGENy scores are computed per cell, and then averaged across barcodes. b) Average expression of genes correlated with diffusion pseudotime for barcodes with missense variants, the barcodes ranked by diffusion score. c) Correlation of JAK-STAT pathway genes with the diffusion score for all barcodes. Significant positive (blue) and negative (red) correlations are highlighted. d) Numbers of gRNAs for the genotyped experiment split by consequence. Several consequences split by “/” indicates that there are several different barcodes for the gRNA with different genotypes and different consequence. e) As d, but performed separately for each predicted consequence. f) Percentage of barcode edits within and outside of the 4-9 base editing window, where position 21-23 is the protospacer adjacent motif (PAM), for barcodes with JAK1 targeting gRNAs. g) Genes regulation differences between SoF and full-impact mutations. The plot shows genes that are upregulated for SoF variants compared to high-score missense and not downregulated compared to benign mutations (high AUC to high-score missense, blue), or downregulated for SoF variants compared to high-score missense and not upregulated compared to benign variants (red).

**Supplementary Table 1 - Information associated with individual cellular barcodes**

Information on the 233 most confidently called cellular barcodes including identity of puroR and iBAR barcodes (puroBC, gRNA_iBAR), called genotype (GT), presence and nature of homozygous edits (any homozygous edits, homozygous edits), targeted gene, position of gRNA in hg38 (start, strand, end, exonic). It also contains details of the actual consequences (consequence of each individual mutation, worst consequence) from variant effect predictor as well as those predicted from gRNA sequence (predicted consequence). Variants present in the COSMIC database (existing variation) or other databases (citation) are also noted. We show the functional effect of the variant using FACS score and proliferation score^8^, the diffusion score (this study), PROGENy pathway score. Information on the guide RNA sequence (gRNA_seq), whether the edit was within the expected editing window (editing positions and out_of_window_edit) and predicted and actual edited sequences (predicted_seq_edited or actual_edit_hom) and whether these were the same (actual_equal_predicted_editing) are also shown.

**Supplementary Table 2 - Primer sequences**

The primer sequences (5’-3’) used for construction of the gRNA library (library construction), detection of the barcodes in scRNAseq (single cell barcode detection), for genotyping of the JAK1 gene (JAK1 amplicon panel) and the partial pKLV sequence are indicated.

